# Biofoundry-assisted Golden Gate cloning with AssemblyTron

**DOI:** 10.1101/2023.11.28.569037

**Authors:** John A. Bryant, R. Clay Wright

## Abstract

Golden Gate assembly is a requisite method in synthetic biology that facilitates critical conventions such as genetic part abstraction and rapid prototyping. However, compared to robotic implementation, manual Golden Gate implementation is cumbersome, error-prone, and inconsistent for complex assembly designs. AssemblyTron is an open-source python package that provides an affordable automation solution using open-source Opentrons OT-2 lab robots. Automating Golden Gate assembly with AssemblyTron can reduce failure-rate, resource consumption, and training requirements for building complex DNA constructs, as well as indexed and combinatorial libraries. Here, we dissect a panel of upgrades to AssemblyTron’s Golden Gate assembly capabilities, which include Golden Gate assembly into modular cloning part vectors, error-prone PCR combinatorial mutant library assembly, and modular cloning indexed plasmid library assembly. These upgrades enable a broad pool of users with varying levels of experience to readily implement advanced Golden Gate applications using low-cost, open-source lab robotics.

## 1. Introduction

Golden Gate assembly is a principal enabling technology in synthetic biology, and automated implementation has transformed it into a high throughput genetic engineering tool. Golden Gate assembly is a scarless DNA assembly strategy capable of assembling up to 40 fragments (*1–4*). Plasmid libraries of standardized modular parts are often used to facilitate rapid Golden Gate assembly, circumventing PCR amplification. Originally, the Modular Cloning system (MoClo) implemented this strategy and has inspired a wide array of plasmid toolkits for different applications (*5–9*), which have been reviewed by Casini et al. (*10*). Modular cloning toolkits define a standard of unique and seamless overhangs for predefined part types, which are then used to build libraries of genetic parts such as promoters, terminators, markers, coding sequences, etc that can be assembled in combinatorial fashion. In the Sky biofoundry, RoboMoClo combines lab automation with the MoClo system to create a biofoundry-assisted Golden Gate platform for metabolic pathway-level DNA assembly (*11*). However, RoboMoClo does not provide openly available code, requires expensive robotics, and is intended as a service platform rather than a transferable tool.

Prior to AssemblyTron, resources for automating customized, in-lab Golden Gate workflows were scarce. Golden Gate assembly is often performed with commercial kits, where just one or two fragments are inserted into a predefined entry vector – a gross underutilization of the method. Customized Golden Gate greatly expands the flexibility and potential construct complexity, however manual Golden Gate implementation requires careful design and extensive molecular biology background knowledge (*3*). Fortunately, many automation tools exist for designing custom Golden Gate assemblies (*12–15*), but manually interpreting and implementing their output files is labor intensive and error-prone. Modular cloning toolkits, such as the MoClo system, can simplify Golden Gate workflows, however PCR is unavoidable for integrating a gene of interest into a part plasmid. Moreover, the complex, multi-level nature of toolkits limits their accessibility for inexperienced researchers. We designed AssemblyTron to help resolve these issues with automation.

AssemblyTron is an open-source python package that, when paired with an Opentrons OT-2 lab robot, can provide biofoundry capabilities with minimal startup investment (*16*). AssemblyTron 1.0 uses j5 DNA design files as input and generates fully customized OpenTrons Application Program Interface (API) protocol scripts for assembling plasmids via homology mediated assembly or Golden Gate. We previously validated that AssemblyTron is adept at automating multi-part Golden Gate assembly and can simultaneously increase throughput, eliminate human errors, and reduce training demands (*16*). AssemblyTron adds to a very sparse field of open-source build tools for automating Golden Gate assembly. Since our initial publication, we have expanded AssemblyTron’s toolkit of capabilities.

Here, we will review AssemblyTron 2.0 upgrades for automating biofoundry-enabling Golden Gate protocols. First we describe a modified version of AssemblyTron’s standard Golden Gate assembly protocol, which accepts any Type IIS restriction-enzyme-digestible part entry vector as the construct backbone. Next, we introduce a protocol for assembling mutant libraries with Golden Gate. In this strategy, AssemblyTron uses error-prone PCR with tunable error-rates to mutate user-specified DNA fragments that are assembled into plasmid libraries via Golden Gate. Finally, we introduce the modular cloning toolkit builder, which is the most recent and developmental installment of AssemblyTron. This feature provides an interactive graphical user interface (GUI), inspired by Figure 2 of the Yeast MoClo Toolkit publication (*7*). The current toolkit builder offered by AssemblyTron is for the auxin modular cloning toolkit, which is a custom, in-house toolkit in the Wright Lab (*see* **Note 1**). However, our automation strategy for toolkit building can be adapted for any toolkit system as a fork or pull-request of the AssemblyTron github repository https://github.com/PlantSynBioLab/AssemblyTron.

## 2. Materials

### 2.1 Hardware

1. Opentrons OT-2 liquid handling robot, equipped with temperature module and thermocycler module (*see* **Note 2**).
2. Biorad thermocycler, or any gradient thermocycler.
3. Nanodrop-2000c Microvolume Spectrophotometer (Thermo Fisher) or similar.
4. Agarose DNA gel electrophoresis setup.
5. iBright imaging system (Thermo Fisher), or similar, for imaging DNA bands.

### 2.2 Software

1. AssemblyTron (https://github.com/PlantSynBioLab/AssemblyTron) (*16*)
2. Python (https://www.python.org/downloads/)
3. J5 and DIVA (https://j5.jbei.org/), for plasmid design (*12*).
4. Anaconda (https://www.anaconda.com/download), for managing AssemblyTron software dependencies (OS, Pandas, Shutil, Numpy, Subprocess, Datetime, Tkinter, CSV, and Opentrons).
5. OpenTrons run app (https://opentrons.com/ot-app/)

### 2.3 PCR

1. Phusion High-Fidelity DNA Polymerase (NEB), supplied with 10x buffer, or any other high-fidelity polymerase, for amplification of backbone fragments.
2. Taq Polymerase (NEB), supplied with 10x magnesium-free buffer and 25 mM MgSO4, for error-prone fragment amplification.
3. Custom primers can be ordered from any commercial vendor.
4. Monarch Plasmid MiniPrep kit (NEB) for isolating template plasmid and assembled plasmid from E. coli.
5. Monarch DNA Gel Extraction kit (NEB), for purifying PCR products products prior to assembly (Note: protocol specifies when to use Gel extraction or DNA clean and Concentrate kit)
6. DNA Clean and Concentrate kit (Zymo Research), for purifying PCR products prior to assembly (Note: protocol specifies when to use Gel extraction or DNA clean and Concentrate kit)
7. DNA ladder: 1 kb DNA Ladder (NEB) for gel electrophoresis
8. 50x TAE buffer, prepared according to Green et al., 2012 (*17*).
9. GelRed® Nucleic Acid Gel Stain (Biotium)
10. For preparing gels for electrophoresis: 0.5% agarose in 1x TAE is melted in a microwave oven, and 3 μL of GelRed® stain is added per 50 mL of agarose.
11. Running buffer for agarose gels is 1xTAE.

### 2.4 Cloning

1. Template plasmid Genbank files for plasmid design.
2. Restriction endonuclease BsaI (10 U/μL) (NEB), supplied with 10x NEB Buffer.
3. T4 DNA Ligase (2,000 U/μL) (NEB), supplied with 10x T4 DNA Ligase Reaction Buffer.
4. BSA, Molecular Biology Grade (NEB), or Recombinant Albumin, Molecular Biology Grade (NEB), diluted to 10 mg/mL.
5. Lysogeny Broth (LB), Miller (Fisher BioReagents). 1% (w/v) bactoagar was added for plates.
6. Antibiotics: ampicillin and kanamycin. Filter-sterilized stocks of 100 mg/mL ampicillin and 50 mg/mL kanamycin (stored in aliquots at -20 C) are diluted 1:1,000 (final concentration: 100 ug/mL ampicillin and 50 ug/mL kanamycin) in autoclaved and cooled liquid medium.
7. High efficiency competent *E. coli* cells (Top10 chemically competent *E. coli* for standard plasmid assembly and NEB10B electrocompetent *E. coli* for library assembly.

### 2.5 Sequencing and Screening

1. Plasmid assembly can be assessed by chromoprotein expression, Sanger sequencing aligned with A Plasmid Editor (*18*), and whole plasmid sequencing.
2. Custom sequencing primers can be design for any construct using A Plasmid Editor (*18*)

## 3. Methods

AssemblyTron was developed and debugged using Windows OS, so we recommend implementing it with Windows OS. (*see* **Note 3**). The most up to date version of AssemblyTron is available at https://github.com/PlantSynBioLab/AssemblyTron. Prior to installing AssemblyTron, users need the latest versions of Python and R installed on their OT-2 connected computers. The package can also be downloaded from pypi. We recommend using Anaconda to easily install and manage AssemblyTron dependencies, which are listed in section 2.2. The OpenTrons run app should also be installed for uploading AssemblyTron scripts to the OT-2.

### 3.1 Plasmid Design and preliminary setup

A plasmid design must be prepared with j5 and DIVA prior to building it with AssemblyTron. For entry vector backbone parts, the forced assembly strategy should be set to “digest.” Eugene rules should be added to specify which parts belong together, unless a complete combinatorial design is desired. When running the j5 algorithm, the design strategy should be set as “Combinatorial Golden Gate.” (*see* **Note 4**). After running j5, download results, unzip the compressed file, and store the design folder on your local hard drive. Primers can be found in `masteroligolist.csv` or in the `pmas00001_combinatorial.csv` design file and must be ordered prior to moving forward. To ensure successful PCR amplification, digest any circular template plasmids with a single-cutter restriction endonuclease prior to PCR amplification (*see* **Note 5**). Optionally, entry vectors can be digested with type IIS restriction endonuclease and gel purified to reduce template vector carry-through. This step can be skipped if the entry vector contains a toxicity gene (e.g. ccdB) or a fluorescent marker that distinguishes it from positive transformants.

### 3.2 Entry Vector Golden Gate Assembly

AssemblyTron enables flexible Golden Gate assembly into any entry vector, which is a critical capability for assembling level 0 part plasmids for modular toolkits as well as reducing dependency on commercial Golden Gate kits.

1. Navigate to the `∼/AssemblyTron/src/AssemblyTron` directory on your local drive. Go to the `Golden_Gate/` folder, and select the `Setup_digests_gradient.py` script. (*see* **Note 6**).
2. Use the dialog box to navigate to your j5 design folder, which was previously saved to the local hard drive (section 3.1). Select confirm, and AssemblyTron automatically parses the j5 design file by importing a parsing module (*see* **Note 7**).
3. Screen the `reagent_setup.txt` file, which will be used to stage primers, templates, reagents, labware, and tubes on the OT-2 deck. Ensure that all templates and primers are assigned to a deck position. Close it until needed.
4. A parameter selection dialog box with default values for PCR and assembly is displayed. Most defaults do not need to be modified, however the following parameters will require consideration or adjustment:
  a. Check the Gradient PCR or touchdown PCR option. The default is touchdown PCR, but this may need to be changed depending on the desired DNA yield (*see* **Note 8**) or how prone the PCR is to failure (e.g., if the range of annealing temperatures for fragments is large, it is safer to use the gradient amplification option.)
  b. Check the “PaqCI?” option. This specifies whether PaqCI or a different type IIS restriction enzyme is being used for Golden Gate assembly. PaqCI requires an activator additive, which alters reagent volumes in the final Golden Gate assembly mixes.
  c. There are two blank parameter entry windows available so that users can add additional parameters to protocols without modifying the dialog box generator scripts. These do not have to be entered.
  d. Entry windows for template concentrations (ng/μL) must be entered to proceed.
5. Select confirm once parameters are updated.
6. A dialog box is displayed for entering concentrations of the entry vector backbones used in this assembly. Enter concentrations in ng/μL and select confirm. (*see* **Note 9**).
7. In the next dialog box, select which parts of the protocol to run. Users can choose from Primer and template dilution, PCR preparation, DpnI digest (template removal), Golden Gate assembly setup, and Golden Gate assembly run. This option makes it easy to skip certain parts of the protocol without modifying python code. For example, primers may already be diluted, so that part of the protocol would not need to be selected.
8. Select confirm. The gradient optimization algorithm runs automatically, unless the touchdown PCR option was specified.
9. Screen the `reaction_setup.txt` file, which specifies where to place tubes for PCRs and final assemblies. Ensure that each PCR and assembly is assigned to a tube and close the file until needed.
10. The setup script imports the primer dilution and Golden Gate protocol writer modules (`dilution_24_digests_writer.py` and `GoldenGate_digests_separatepcrruns_gradient_writer.py`) and executes them automatically. The final customized protocol scripts then open automatically. Close them to allow the setup script to finish running.
11. Navigate to the new directory, labeled with the current date, that was just generated.
12. Protocol scripts are ready to be uploaded directly to the OT-2. (*see* **Note 10**).
13. See section 3.5 for instructions on uploading protocols to the OT-2.
14. Amplify fragments, run Golden Gate assembly, clean and concentrate assembled plasmids, and transform into high efficiency chemically competent *E. coli*.

### 3.3 Mutant Library Assembly

Mutant library screening is a common strategy for protein engineering and directed evolution (reviewed by Wang et al.) (*19*). AssemblyTron provides protocols for facilitating mutant library assembly with error-prone PCR and Golden Gate assembly. Targeted mutagenesis of gene regions followed by assembly with unaltered backbone fragments provides high-quality plasmid libraries for functional screening (*20, 21*). Using AssemblyTron to automate library assembly ensures consistent library quality, which saves time and resources. AssemblyTron allows users to tune error rate (Figure 1A), which accommodates various target fragment sizes and libraries of varying degeneracy. We assembled libraries of pink chromoprotein expression plasmids with different error rates and screened for functional variation by colony color (Figure 1B, 1C). Our data indicates that increasing error-rate settings with AssemblyTron correlate with increasing loss-of-function colorless colonies and diminishing pink colonies.

**Figure 1:**
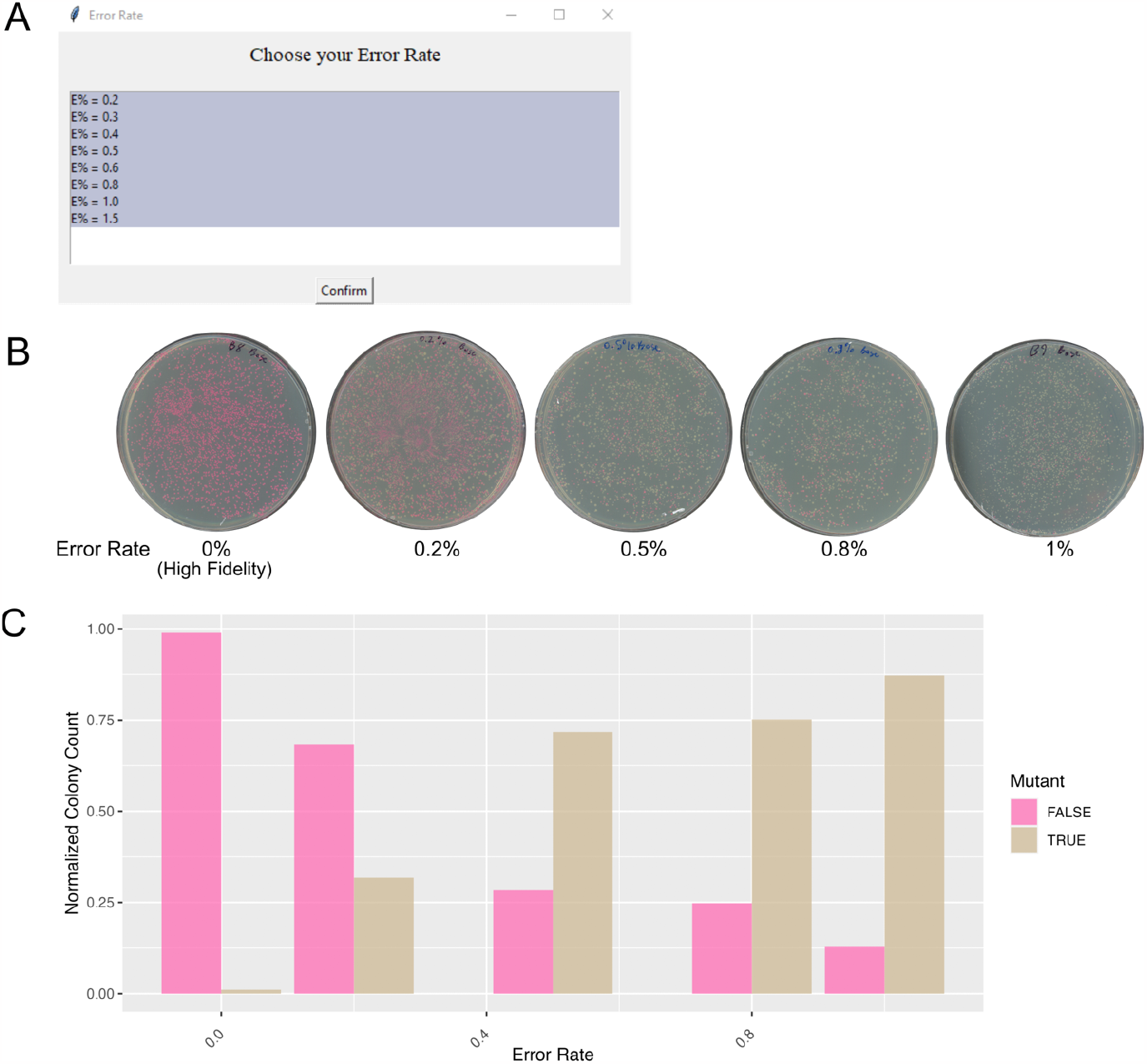
Tuning mutagenesis with AssemblyTron error prone PCR. A) A dialog box provided by AssemblyTron is used to tune the error rate. B) Qualitative mutant accumulation increases as higher mutation rates are specified with AssemblyTron. C) Quantitative mutant accumulation increases as higher mutation rates are specified based on a plot of normalized colony count for 10X dilutions of plates in B.

Here, we provide a detailed protocol for using AssemblyTron’s library assembly feature. Our workflow consists of three stages. In stage one, an initial PCR is performed to amplify backbone fragments and a template amplicon for the error-prone fragment with high fidelity polymerase (*see* **Note 11**). In stage two, error-prone PCR is used to mutate the target fragment with a user-specified error rate (Figure 1A). In stage three, Golden Gate is used to assemble the mutant fragment pool with backbone fragments.

Golden Gate Primers with 5’ Golden Gate overhangs (*see* **Note 12**), should be ordered prior to implementing this protocol. Additionally, shorter primers lacking 5’ Golden Gate overhangs must be ordered to generate the initial template amplicon in stage one. This template will be used in stage two error-prone PCR. Primers lacking Golden Gate overhangs prevent the initial template amplicon from being assembled into the library in stage three, which would decrease the overall error-rate of the library.

*Stage 1*

1. Navigate to the `∼/AssemblyTron/src/AssemblyTron/` directory and go to the `Error_prone_PCR_Golden_Gate/` folder.
2. Select a stage one setup script. `1_Setup_STD_GoldenGate_digests_PCR.py` is for libraries that will be assembled into entry vector backbones, and `1_Setup_STD_GoldenGate_PCR.py` is for libraries that will be assembled strictly from PCR amplicons. (*see* **Note 13**). Click on the appropriate script.
3. Use the dialog box to navigate to your j5 design folder, which was previously saved to the local hard drive (section 3.1). Select confirm, and AssemblyTron automatically parses the j5 design file by importing a parsing module (*see* **Note 7**).
4. Screen the reagent_setup.txt file, which will be used to stage primers, templates, reagents, labware, and tubes on the OT-2 deck. Ensure that all templates and primers are assigned to a deck position. Close it until needed. (*see* **Note 14**).
5. A parameter selection dialog box with default values for PCR and assembly is displayed. Most defaults do not need to be modified, however the following parameters will require consideration or adjustment:
  a. Check the Gradient PCR or touchdown PCR option. Stage one PCR is set to the gradient option by default. This is to ensure clean bands with high DNA concentration for high library quality and successful yields after gel extraction. (*see* **Note 15**).
  b. Template concentrations (ng/μL) must be entered to proceed.
6. Select confirm once parameters are updated.
7. In the next dialog box, select which parts of the protocol to run. Users can choose from Dilution and PCR mix in this stage, since assembly will not be performed until stage three. (*see* **Note 16**).
8. Screen the reaction_setup.txt file, which specifies where to place PCR tubes on the on-deck thermocycler. Ensure that each PCR is assigned to a tube and close the file until needed.
9. The setup script imports the primer dilution and Golden Gate protocol writer modules (`dilution_24_digests_writer.py` and `GoldenGate_digests_separatepcrruns_gradient_writer.py`) and executes them automatically. The final customized protocol scripts open automatically. Close them to allow the setup script to finish running.
10. Navigate to the New dated folder (e.g., `202310011038_PCR/`).
11. The final customized protocol scripts that are ready to be uploaded directly to the OT-2. (*see* **Note 10**). See section 3.5 for instructions on uploading protocols.
12. After initial dilution and PCR protocols run, isolate fragments with gel electrophoresis, extract fragments, and measure DNA concentration before proceeding to stage two. (*see* **Note 17**). *Stage 2*
13. Navigate to the `∼/AssemblyTron/src/AssemblyTron/Error_prone_PCR_Golden_Gate/` directory and go to the dated PCR folder generated in stage one (e.g., `2023010011038_PCR/`).
14. Click on the stage two setup file (e.g., `2_2023010011038_Setup_Error_prone_PCR.py`)
15. Use the dialog box to navigate to your j5 design folder, which was previously saved to the local hard drive (section 3.1). Select confirm, and AssemblyTron automatically parses the j5 design file by importing a parsing module (*see* **Note 7**).
16. Use the next dialog box to select which fragments in the design are targets for mutagenesis by placing a 1 in the entry box. This specifies which fragments will be amplified with error-prone PCR.
17. Screen the `reagent_setup.txt` file, which will be used to stage primers, templates, reagents, labware, and tubes on the OT-2 deck. Ensure that all templates and primers are assigned to a deck position. Close it until needed.
18. A parameter selection dialog box with default values for PCR is displayed. Template concentrations are the only entries necessary, and defaults do not need to be modified. (*see* **Note 18**).
19. In the next dialog box, select which parts of the protocol to run. Users can choose from Dilution and PCR mix in this stage, since assembly will not be performed until stage three (*see* **Note 19**).
20. Choose an error rate in the next dialog box (Figure 1A). Depending on the user-specified error rate, AssemblyTron will alter ratios of manganese to magnesium and dNTPs to tune error rate and minimize transition/transversion bias (Figure 1B) (*see* **Note 20**).
21. The setup script imports primer dilution and error-prone PCR protocol writer modules (`dilution_Error_prone_PCR_writer.py` and `Error_prone_PCR_writer.py`) and executes them automatically. The final customized protocol scripts open automatically. Close them to allow the setup script to finish running.
22. Navigate to the new stage two folder that was just created (e.g., `2_202310021038_EPPCR/`).
23. The final customized protocol scripts that are ready to be uploaded directly to the OT-2. See section 3.5 for details on uploading protocols to the OT-2. (*see* **Note 10**).
24. Run error-prone PCR on OT-2, clean and concentrate, and measure concentration of PCR products before proceeding to stage three. *Stage 3*
25. Navigate to the `∼/AssemblyTron/src/AssemblyTron/Error_prone_PCR_Golden_Gate/` directory and go to the dated PCR folder generated in stage one (e.g., `2023010011038_PCR/`).
26. Click on the stage three setup file (e.g., `3_2023010011038_Assembly.py`)
27. Use the dialog box to navigate to your j5 design folder, which was previously saved to the local hard drive (section 3.1). Select confirm, and AssemblyTron automatically parses the j5 design file by importing a parsing module (*see* **Note 7**).
28. Screen the `reagent_setup.txt` file, which will be used to stage reagents and labware on the OT-2 deck. Close it until needed.
29. A parameter selection dialog box with default values for assembly is displayed. Most defaults do not need to be modified, however the following parameters will require consideration or adjustment:
  a. Check the “PaqCI?” option. This specifies whether PaqCI or a different type IIS restriction enzyme is being used for Golden Gate assembly. PaqCI requires an activator additive, which alters reagent volumes in the final Golden Gate assembly mixes.
  Select confirm.
30. In the next dialog box, select which parts of the protocol to run. Users can choose from Golden Gate setup and Golden Gate run in stage three, since PCR steps were performed in stages one and two.
31. Use the next dialog box for uploading concentrations of each assembly fragment in ng/μL. Each entry box is labeled by well number to avoid confusion. A new window will appear for each assembly (e.g., if there are four assemblies, four concentration input windows will appear consecutively).
32. Screen the `reaction_setup.txt` file, which specifies where to place fragment PCR tubes and final assembly PCR tubes on the on-deck thermocycler. Close the file until needed for setting up the OT-2 deck.
33. The setup script imports the Golden Gate protocol writer module (`GoldenGate_digests_separatepcrruns_gradient_writer.py`) and executes it automatically. The final customized protocol script opens automatically. Close it to allow the setup script to finish running.
34. Navigate to the new stage three folder that was just created (e.g., `3_202310021038_Assembly/`).
35. The final customized protocol script that is ready to be uploaded directly to the OT-2. (*see* **Note 10**). See section 3.5 for details on uploading protocols to the OT-2.
36. Run the library assembly on the OT-2, clean and concentrate, and transform into high efficiency competent *E. coli* cells.

### 3.4 Modular Cloning toolkit builder

Automated modular cloning indexed plasmid library assembly has proved to be a successful platform strategy for biofoundries (*11*). Combinatorial part assembly and shuffling will inevitably remain a useful cloning strategy even as DNA synthesis prices become trivial, so we have begun offering it as an AssemblyTron feature. Here we introduce a preliminary, open-source protocol for automating construct assembly from an auxin modular toolkit we have built in-house, based on part standards of the Yeast Modular Cloning Toolkit (*7*). This toolkit build strategy will serve as a model for expanding AssemblyTron platform to other modular cloning toolkits in the future.

1. Navigate to the `∼/AssemblyTron/src/AssemblyTron/` directory and go to the `MoClo_builder/` folder.
2. Select the `Setup_MoClo.py` script by clicking on it.
3. Use the dialog box to specify how many construct variants you are assembling from the toolkit.
4. A graphical user interface for selecting modular parts appears (Figure 2). Choose the desired combination of parts. One part from each column must be selected, for a total of eight (or more, *see* **Note 21**) parts (Figure 2A, 2B).
5. Select confirm after selecting parts for the first construct. New selection windows will be consecutively displayed for specifying each additional construct.
6. A parameter selection dialog box with default values for assembly is displayed. Most defaults do not need to be modified. Select confirm.
7. Navigate to the new dated folder that was just created (e.g., `202310021254_MoClo/`).
8. Run the `MoClo_writer.py` protocol writer script by clicking on it. This will generate the final customized protocol script that is ready to be uploaded directly to the OT-2.
9. See section 3.5 for details on uploading protocols to the OT-2.
10. Run construct assembly on the OT-2, clean and concentrate, and transform into high efficiency chemically competent E. coli.

**Figure 2:**
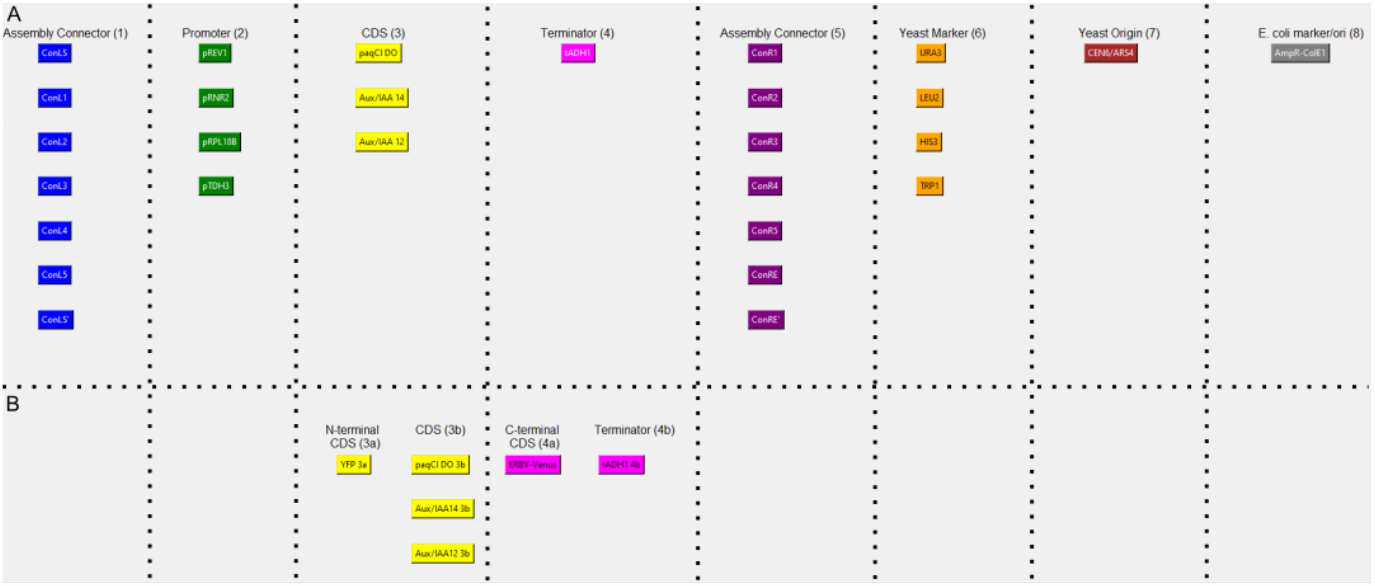
AssemblyTron graphical user interface for modular part selection. A) Any combination of eight color-coded part types can be specified for assembly. B) A pair of type 3a and 3b parts can be selected in place of a standard type 3 part for N-terminal protein fusions. Similarly, a type 4a and 4b part can be selected in place of a standard type 4 part for a C-terminal protein fusion.

### 3.5 Uploading and running protocols on the OT-2

1. Open the Opentrons run app, turn on the OT-2 and modules, and connect the app to the OT-2.
2. Navigate to the protocols tab on the left side of the app, select import, and upload protocol scripts in the correct order.
3. Open `reagent_setup.txt` and `reaction_setup.txt`. Use these files to guide physical setup of the opentrons deck (*see* **Note 22, 23**).
4. Click start run. Facilitate protocols by following any directions that are displayed on the OpenTrons run app during protocol pauses.

### 4. Notes

1. The auxin modular cloning toolkit should be published by the end of 2024.
2. Minor modifications could be made to the AssemblyTron source code to remove dependency on the modules and avoid purchasing them.
3. While running AssemblyTron on Mac OS may be possible, it will likely require additional debugging and programming knowledge.
4. Update “Golden Gate Recognition sequence” and “Golden Gate Term Extra Seq” fields within the “j5 parameters” if a different type IIS restriction enzyme besides the default BsaI is being used.
5. Digesting circular template plasmids with a single-cutter restriction endonuclease prior to PCR amplification is optional, but skipping this step often leads to PCR failure.
6. If the design does not use an entry vector, select an alternative setup script. For standard Golden Gate with 24 primers and templates, select `Setup_nodigests_seppcr_gradient_24.py`. For scaled standard Golden Gate with 96 primers and templates, select `Setup_nodigests_seppcr_gradient_96.py`. The workflow for each of these is nearly identical to Entry Vector Golden Gate.
7. The j5 parsing modules are located at `∼/AssemblyTron/src/AssemblyTron/`. Either `j5_to_csvs.py` or `j5_to_csvs+digests.py` is used depending on the assembly design.
8. Gradient amplification typically yields cleaner bands and higher DNA yields, while touchdown amplification has a higher likelihood of producing nonspecific bands and lower DNA yields.
9. In standard Golden Gate assembly, this dialog box does not appear.
10. A batch transfer script is no longer necessary for using AssemblyTron, thanks to the protocol writer modules.
11. We recommend gel purifying this template amplicon to reduce template DNA carry-through to the final library. This increases library quality.
12. We typically use j5/DIVA, https://public-diva.jbei.org/, to design these primers (*12*).
13. We recommend assembling libraries into entry vectors, since this typically increases assembly efficiency by reducing the number of fragments in the assembly. This also prevents unnecessary backbone PCR amplification.
14. Use the shortened 3’ only primers lacking Golden Gate overhangs for the initial template amplicon to be used in stage 2.
15. Touchdown PCR isn’t a good option for error-prone PCR stage 1 since it is more likely to amplify off-target fragments and sometimes doesn’t provide the highest possible concentration of amplified product.
16. There is no need for methylated templated digestion with DpnI in the error-prone PCR workflow since all fragments are gel extracted.
17. Gel purification reduces template carry-through and ensures a high quality library at the conclusion of the workflow.
18. Remember to use the amplicon from stage 1 as template for error-prone PCRs in stage 2.
19. There is no need for methylated templated digestion with DpnI in the error-prone PCR workflow since mutant fragments are amplified from a PCR amplicon with no Golden Gate overhangs.
20. PCR tubes for error-prone amplification should be placed in the same location on the thermocycler as their initial template amplicons from stage 1.
21. A combination of a type 3a and a type 3b part may be selected in place of a standard type 3 part to make an N-terminal protein fusion. Similarly, a type 4a and a 4b part combination may be selected in place of a standard type 4 part to make a C-terminal protein fusion.
22. The Opentrons run app provides a deck map under labware setup>map view.
23. Ensure that shortened 3’ only primers lacking Golden Gate overhangs are used to generate the initial template amplicons in error-prone PCR stage one. Backbone fragments should be amplified from standard Golden Gate overhang primers in stage one.

## Acknowledgements

We thank Cameron Longmire, Mason Kellinger, Sriya Sridhar, Ryan Miller and Samuel Janousek for their contributions to the AssemblyTron package.

## Notes

### Competing Interest Statement

The authors have declared no competing interest.

